# Regulation of gene expression under high hydrostatic pressure: the versatile role of the master regulator SurR in energy metabolism

**DOI:** 10.1101/2025.03.14.643387

**Authors:** Yann Moalic, Toan Bao Hung Ngyuen, Jordan Hartunians, Tiphaine Birien, Axel Thiel, Mohamed Jebbar

**Affiliations:** Univ Brest, Ifremer, CNRS, EMR 6002 BIOMEX, BEEP, F-29280 Plouzané, France

**Keywords:** Piezophilic archaea, High hydrostatic pressure (HHP), Energy metabolism, Transcriptional regulation, Sulfur metabolism, Gene expression, Adaptation to extreme environments

## Abstract

In *Thermococcus barophilus*, a piezophilic hyperthermophilic archaeon, the expression of several gene clusters, including those of energy metabolism, is modulated by hydrostatic pressure. In *Thermococcales*, SurR, a redox-sensitive transcriptional regulator that responds to sulfur availability, regulates genes involved in energy metabolism. To better understand how high hydrostatic pressure (HHP) influences the expression of energy metabolism genes, several gene deletion mutants including *surR* partial knockout, were constructed and analyzed under various culture conditions, including different hydrostatic pressures and the presence or absence of sulfur.

Phenotypic analysis of the *surR* mutant revealed that SurR affects both growth and gene expression, independently of sulfur availability. This regulatory behavior differs from that observed in non-piezophilic Thermococcales species such as *Pyrococcus furiosus* and *Thermococcus kodakarensis*. These findings suggest that hydrostatic pressure influences the physiological role or functional state of SurR in *T. barophilus*, highlighting its adaptive versatility in extreme environments.

**IMPORTANCE:** This study provides new insights into the adaptive mechanisms of hyperthermophilic archaea to high hydrostatic pressure, a key factor in deep-sea environments. By demonstrating that SurR regulation differs in *T. barophilus* compared to non-piezophilic species, it suggests that pressure can modify transcriptional control mechanisms, potentially reshaping energy metabolism strategies in deep-sea archaea. Understanding these regulatory adaptations contributes to our broader knowledge of microbial life under extreme conditions and may have implications for biotechnology, particularly in designing pressure-resistant enzymes or metabolic pathways.

## INTRODUCTION

*Thermococcales* are ubiquitous extremophilic Archaea found at hydrothermal environments growing optimally at temperature over 80°C (Schut et al., 2014). The chronic energy stress induced by high temperature requires specific metabolic strategies allowing them to thrive in these environments (Valentine, 2007). Additionally, *Thermococcales* can oxidize different types of carbon-based molecules (proteins, organic acids, carbohydrates…) to produce energy (Nakagawa and Takai, 2006).

A diverse set of hydrogenases, oxidoreductases, and electron transporters enable *Thermococcales* to generate the electron fluxes for maintaining the ionic gradient across the membrane, which is directly linked to ATP production. Consequently, energy conservation is associated with both H_2_S production and H_2_ turnover and recycling (Wu et al., 2018). While some aspects of the functional mechanisms of these catabolic components remain unclear, they are relatively well characterized. The membrane-bound hydrogenase (MBH) and its homologous S^0^-reducing reductase (MBS) are conserved across all publicly available *Thermococcales* genomes (Schut et al., 2013). These enzymes oxidize the ferredoxin yielding either H_2_S or H_2_, while contributing to energy conservation via Na+ pumping. In *P. furiosus*, one of the most study model of *Thermococcales* with *T. kodakarensis,* the addition of elemental sulfur (S^0^) to growth medium, induces the expression of MBS-encoding genes, while repressing MBH-encoding genes (Chou et al., 2007). This shift results in a two-fold increase in microbial cell yield (Schut et al., 2007), suggesting that MBS is more efficient in energy conservation. MBS facilitates electron transfer for the reduction of polysulfide chains while simultaneous driving Na^+^ translocation across the membrane. Additionally, it should be noted that in the presence of S^0^ H_2_S can be produced abiotically (Wu et al., 2018;Burkhart et al., 2019;Yu et al., 2020).

The master redox-active transcription regulator, SurR, modulates the expression of various enzymes of the energy conservation system based upon whether elemental sulfur is available or not (Lipscomb et al., 2009). When sulfur (S^0^) is present, SurR becomes oxidized and no longer binds specifically to its target DNA. As a result, the H_2_-related genes are deactivated, while the H_2_S-related genes are de-repressed, prompting *P. furiosus* to produce hydrogen sulfide (Yang et al., 2010). In absence of sulfur, the reduced form of SurR represses genes implicated in H_2_S production while activating those linked in H_2_ production, leading *P. furiosus* to produce hydrogen.

The only available SurR structure was solved for the *P. furiosus* variant (Yang et al., 2010). SurR is a homodimer with double symmetry, where each monomer consists of three distinct domains. The N-terminal (residues 1–75) and C-terminal (residues 166–219) regions each form a winged helix-turn-helix (wHTH) DNA-binding fold related to that of the ArsR family (Yang et al., 2010).

The SurR recognition motifs (GTTn3ATC or GTTn3AACn5GTT) are located in many promoters of genes involved in the sulfur response of *Thermococcales* (Lipscomb et al., 2009;Hidese et al., 2017;Lim et al., 2017;Lipscomb et al., 2017;Moalic et al., 2021). A redox-active switch controls DNA binding, through a CxxC motif in the N-terminal wHTH domain, likely by affecting the conformation of the protein’s DNA binding domain. In the reduced free thiol form, in the absence of S^0^, SurR activates the expression of its own gene as well as genes involved in hydrogen metabolism. SurR also represses the expression of Nsr (NADPH-dependent sulfur reductase), Mbs and other clusters involved in sulfur reduction, such as Pdo (Protein disulfide oxidoreductase) (Lipscomb et al., 2017). However, when S^0^ is supplied, the CxxC motif forms a disulfide bond, abrogating specific DNA binding by SurR, resulting in deactivation of the hydrogenase genes and de-repression of sulfur reducing genes. Indeed, as described in *T. kodakarensis*, *surR* deletion inhibits growth in the absence of sulfur but has no effect in the presence of sulfur (Santangelo et al., 2011). Activation or repression by SurR is linked to its binding site position in the promoter. At target sites located upstream of the TATA box, SurR acts as an activator, whereas it represses transcription when it binds downstream of the TATA box (Yang et al., 2010). In some cases, the presence of both upstream and downstream binding sites is essential for repression (Hidese et al., 2017).

An *in vitro* study on *Thermococcus onnurineus*, a member of the Thermococcales, demonstrated that Pdo (protein disulfide reductase) facilitates electron transfer between thioredoxin reductase (TrxR) and the redox-sensitive transcription factor SurR (Lim et al., 2017). These findings indicate that the TrxR-Pdo pair functions as a redox system that reduces SurR. While this reduction has only been observed *in vitro*, the results are significant, as they suggest that SurR-mediated regulation of the S₀ response could be reversed even in the absence of elemental sulfur (S₀) (Lim et al., 2017). Notably, the genomic organization of *surR* is highly conserved across all known Thermococcales genomes. Specifically, *surR* and *pdo* are positioned adjacent to each other in a divergent arrangement, implying that their functions are tightly coordinated *in vivo*. Furthermore, *pdo* has been identified as part of the SurR regulon (Lipscomb et al., 2009). Microarray expression profiling revealed that *pdo* is upregulated during the initial response to S₀ (Schut et al., 2007) and repressed by SurR, which binds to the *pdo* promoter in the absence of S₀ (Lipscomb et al., 2009). It has been proposed that once oxidized sulfur species are depleted within the cell, SurR would revert to its reduced, active state. However, the precise mechanism underlying this transition remains undemonstrated (Yang et al., 2010).

*Thermococcus barophilus* MP, the proposed biological model, is the first true piezophilic hyperthermophile to be isolated from a deep-sea hydrothermal vent (Marteinsson et al., 1999). This archaeon thrives across a wide hydrostatic pressure range of 0.1-80 MPa (P_opt_ 40 MPa) and a temperature range of 48-90°C (T_opt_ 85°C). Elemental sulfur enhances its growth. The complete genome of *T. barophilus* MP has been sequenced and annotated (Vannier et al., 2011) and a genetic system has been developed to further investigate its adaptations to HHP (Thiel et al., 2014;Birien et al., 2018). Neutron scattering studies have revealed that the proteome of *T. barophilus* MP exhibits greater flexibility and sensitivity to HHP, a smaller hydration shell, and more confined intracellular water compared to that of a related piezosensitive species *T. kodakarensis* (Martinez et al., 2016). Interestingly, increased proteome flexibility appears to impair biological function at low pressure, but this effect is mitigated by the accumulation of the osmolyte mannosylglycerate, which stabilizes structures under low pressure conditions (Cario et al., 2016).

Transcriptomic analysis of *T. barophilus* MP has revealed that hydrostatic pressure influences the expression of numerous genes, particularly those involved in hydrogen and elemental sulfur metabolism (Vannier et al., 2015). Notably, the expression of key hydrogenase-encoding genes remains minimal at 40 MPa in the presence of S^0^. However, in a surprising deviation from this pattern, when pressure is reduced to 0.1 or increased to 70 MPa, the expression of the hydrogen-related genes (e.g. *mbh*, *mbh-codh*, *shI* and *shII*) is upregulated by 2 to 40-fold, even in presence of S^0^ (Vannier et al., 2015). In contrast, hydrostatic pressure has little to no effect on the expression of genes within the sulfur regulon, including *mbs*, *nsr*, *surR*, *pdo*, *nfn* and *xfn* in *T. barophilus* MP (Vannier et al., 2015). This suggests that gene regulation in *T. barophilus* differs significantly from that of related non-piezophilic species such as *P. furiosus* or *T. kodakarensis*. The data further imply that high hydrostatic pressure (HHP) may modulate the DNA-binding affinity of SurR in *T. barophilus* MP, contributing to its unique regulatory adaptations to extreme pressure conditions.

In this study, we examined the impact of high hydrostatic pressure (HHP) and the redox regulator SurR on physiological growth and the expression of gene clusters involved in energy conservation. Using a combination of genetic approaches and RT-qPCR, we analyzed growth kinetics and gene expression in various mutants.

Our findings confirm that HHP plays a modulatory role in gene regulation and reveal the existence of a sulfur-independent mechanism influencing SurR activity—an adaptation not observed in non-piezophilic *Thermococcales*. These results underscore a specialized genetic regulatory strategy that enables *T. barophilus* to thrive in deep-sea hydrothermal environments.

## MATERIALS AND METHODS

### STRAINS AND GROWTH CONDITIONS

All the strains used in this study are list in Table S1. A modified version of the Thermococcales Rich Medium (TRM) was used for all growth studies, based on the formulation described by Zeng et al. (2009). This modified medium (TRMm) contains 5 g·L⁻¹ yeast extract and 5 g·L⁻¹ tryptone, an increase from the original TRM composition, to support growth under hydrogenogenic conditions with 5 g·L⁻¹ pyruvate as the carbon source. For sulfidogenic growth, TRMm was supplemented with 0.25 g·L⁻¹ colloidal sulfur, except for induction experiments, where higher sulfur concentrations (0.5 g·L⁻¹ or 2 g·L⁻¹) were used.. Growth experiments were achieved in biological triplicates at 85°C in anaerobic conditions. At atmospheric pressure condition (0.1 MPa), 50 mL vials were filled with 20 mL of medium while cultivation under high hydrostatic pressure (40 MPa and 70 MPa) requires the use of 15 ml vials entirely filled with medium to avoid the imploding risk due to gas phase. The absence of gas phase has no effect on growth at atmospheric pressure (0.1 MPa) comparatively to cultivation realized with a gas phase (data not shown). Each growth kinetic assay was initiated from overnight pre-cultures (16 hours) prepared in the same medium, with an initial cell density of 2 × 10⁶ cells/mL.

For HHP growth experiments, four incubators (Top Industrie) were used (Zeng et al., 2009). Due to compression/decompression constrains, each kinetic point corresponds to a separate incubator containing six 15mL vials, three replicate of the mutant strain and three replicates of the Δ517 control strain (either Δ517 or Δ517p according to the sulfur condition). Growth monitoring was realized by cell counting using a Thoma chamber and photonic microscopy at a magnification of X40.

### MUTANTS CONSTRUCTIONS

The genetic manipulations were realized following the pop-in/pop-out protocol developed in our laboratory (Birien et al., 2018). This genetic tool was developed under sulfidogenic condition, and all the mutant constructions were realized from the Δ517 strain (Table S1): the Mrp-Mbh and Mrp-Mbs membrane clusters (Δ*mbh*, Δ*mbs*), the two cytosolic hydrogenases SHI and SHII (Δ*shI* and Δ*shII*) and the partial deletion of *surR*. More precisely, the double copy of Mrp-Mbh complex was deleted (Δ*MBH* strain) and for the Δ*SurR* strain, a partial deletion was built after all attempts to obtain a completed deletion of its locus have failed. As for *Pyroccocus furiosus* and *Thermococcus kodakarensis*, the 5’ end of the gene containing the CxxC motif was deleted (Santangelo et al., 2011;Lipscomb et al., 2017). Thus, after PCR experiments, some clones were selected and the deletion of the 137bp was confirmed by Sanger sequencing.

All the strains of this study have followed the same protocol of “pyruvate growth adaptation” before testing the effect of sulfur on the phenotypes. The experiments of adaptation were repeated at least 3 times independently with the Δ517 strain as control. Thus, we consider that if the strain cannot growth, it is more probably due to the absence of the gene/gene clusters in the strain than the toxicity of H2-end product, especially when the H2 production pathway is impaired. The growth curves realized in pyruvate condition were carried out with strains that have been 96h into TRMm media before pre-culture preparation.

### GENE EXPRESSION

To assess the effect of SurR regulator on the expression of the cluster of hydrogenases (*mbh1*, *mbh2*, *shI*, *shII*) and sulfane oxidoreductase (*mbs*), RT-qPCR were realized. First, RNA of cells in mid-Log phase were extracted with Trizol reagent. 50 mL of culture were centrifuged at 8000 rpm (6 min at 4°C) and the pellets were suspended in 1 mL of Trizol and transferred in RNase-free 2mL tubes (Biopur Eppendorf). Then the procedural guidelines of the manufacturer were followed until the RNA is suspended in RNase free water (54µL). Then a DNase treatment was realized (RQ1 RNase-free DNase kit, Promega) and the RNA were quantified with a nanodrop 8000 (Thermofischer) and conserved at –80°C. The iScript^TM^ Reverse Transcription Supermix for RT-qPCR (Bio-Rad) was used to get cDNA at the concentration of 2.5 ng.µL^-1^ About 12.5 ng of cDNA were used as matrix for quantitative real-time PCR reactions (SsoAdvanced^TM^ Universal SYBR® Green supermix – Bio-Rad). The primers were used at the final concentration of 0.5µM each in a final volume reaction of 20µL (Listed in Table S3). The reactions were launched on a CFX96TM (Bio-Rad) with an amplification step of 40 cycles (95°, 15s followed by 60°, 30s). The data were saved and analyzed with the Bio-Rad CFX Maestro software.

The ΔΔCt method was then applied to analyze gene expression variation (Schmittgen and Livak, 2008). Due to the diverse growth culture conditions (Sulfur and Pressure), three references genes were tested: two 30S proteins coding (S19 and S13) and the *pcna* (see Table S3 for their Locus).

### GENOME SEQUENCING AND ANALYSIS

DNA was extracted at mid-log growth phase using protocols previously described (Thiel et al., 2014;Birien et al., 2018). Library preparation and Illumina sequencing were performed at Novogene, UK. Bowtie2 was used for read mapping (Langmead and Salzberg, 2012) and Samtools (Li et al., 2009) was used to manipulate and produce the binary data required for the variants detection. Lofreq ((Wilm et al., 2012) is the variant-caller used for inferring single nucleotides variants and indels in the sequencing data comparing to the reference genome of *T. barophilus* MP (Refseq NC_014804.1). Finally, SnpEff helps to predict the effects of genetic variants ((Cingolani et al., 2012)

## RESULTS & DISCUSSION

### Growth of *Thermococcus barophilus* on pyruvate in batch culture

Since its isolation during the French-American ‘MAR93’ cruise (∼30 years ago) (Marteinsson et al., 1999), *T. barophilus* has been routinely cultivated in yeast extract and peptone-based media (YPS and TRM) with added Sulfur for optimal growth (Marteinsson et al., 1995; Zeng et al., 2009). Growth on minimal media (TAA, Thermococcales amino acid or TBM, Thermococcales basic medium) yields lower but interpretable results with sulfur (Thiel et al., 2014; Cario et al., 2015). In rich media without sulfur, growth was minimal or absent due to H₂ toxicity, which inhibits reduced ferredoxin regeneration (Le Guellec et al., 2021) (Malik et al., 1989;Schäfer and Schönheit, 1991). However, continuous culture experiments realized in a gas-lift bioreactor supports high growth rates comparable to sulfur-supplemented batch cultures (Postec et al., 2005). This study required sulfur-free media that did not hinder cell growth for the physiological characterization of energy-conserving gene clusters. After multiple attempts, a modified TRM medium supplemented with pyruvate (5 g/L) successfully supported the growth of *T. barophilus* at satisfactory yields without sulfur. An adaptation period of at least 72h was required for cell growth to reach 6.0E^+07^ cells.mL^-1^ from an initial density of 1.21E^+07^ cells.mL^-1^). However, once adapted, subcultures grew equally well with or without sulfur (FigureS1). This pyruvate-adapted strain originated from the laboratory’s genetic strain, *Tba Δ517* (Birien et al., 2018). The growth rate of Δ517p on pyruvate (0.36 h^-1^) was comparable to that of Δ517 with sulfur and increased slightly to 0.43 h^-1^ when sulfur replaced pyruvate (red curve). In contrast, non-adapted Δ517 exhibited a severe growth impairment (0.004 h^-1^, blue curve). To assess whether genetic mutations c contributed to metabolic adaptation, a genomic comparison was conducted between Tba Δ517p, Tba Δ517 and the reference genome of Tba MP (NC_014804.1). Notably, Tba Δ517 has lost its plasmid pTBMP (NC_15471.1) for unknown reasons, and it is likewise absent in Tba Δ517p. Both strains also lack the gene encoding a phosphoribosyltransferase (TERMP_00517, 648 bp), resulting in a genome assembly of identical size (2,009,585 pb). To identify genetic variants, LoFreq was used to analyze sequence data (Wilm et al., 2012) and categorized potential effects with SNPeff (Cingolani et al., 2012). After filtering for allele Frequency (AF>0.1), *Tba Δ517* exhibited 47 variants compared to Tba MP while Tba Δ517p had 51. Of these, 45 variants in Tba Δ517 were also present in Tba Δ517p, leaving six unique to *Tba Δ517p*. SnpEff analysis identified three variants with a predicted HIGH impact due to frameshift mutations (Table 1). Two of these occur at the same locus (TERMP_02062), which encodes an adenylyl-sulfate kinase (F0LLK7), an enzyme involved in active sulfate synthesis. The third is located at TERMP_00520, annotated as an uncharacterized protein (F0LJW0). Additionally, two mutations were classified as MODERATE impact missense variants. One occurred at locus TERMP_00866 coding for an uncharacterized protein (F0LLR6), causing a Glycine to Arginine substitution at position 159. The other affects TERMP_01561, encoding a low affinity inorganic phosphate transporter (F0LIT4), resulting in a Threonine to Methionine substitution at position 95. Finally, a variant was detected in the promoter region of TERMP_02063, which encodes a transposase. Classified as a MODIFIER impact, this variant likely has uncertain or minimal direct effects, as it does not alter protein sequence. These mutations do not provide clear evidence of direct selection for growth on pyruvate. Instead, the adaptation likely results from metabolic reconfiguration via regulatory mechanisms, similar to shifts observed when sulfur is introduced into pyruvate-grown cells or in comparative proteomic studies of sulfur vs pyruvate grownThermococcales (Schut et al., 2007;Jager et al., 2014;Moon et al., 2015). Future studies are needed to further elucidate these mechanisms. However, this genomic analysis clarified the effects of a 96-hour adaptation period required for growth without sulfur. Thus, we established TRMm + pyruvate as a sulfur-free growth condition for this study.

### SurR binding motif distribution in energy conservation gene promoters

We extended our genomic analysis by systematically screening the *T. barophilus* chromosome for SurR recognition motifs ((GTTn3ATC or GTTn3AACn5GTT). As observed in other Thermococcales, both long and short motifs were detected upstream of genes clusters involved in electron flow (Table2). Notably, *Thermococcales* exhibit diversity in the types and abundance of cytosolic and membrane-bound hydrogenases (Schut et al., 2013). In *T. barophilus*, the membrane-bound hydrogenase exists in two adjascent copies (Mrp-mbh 1 and Mrp-mbh 2, Figure1), (SHI and SHII) whereas the two cytosolic hydrogenases (SHI and SHII) are separated by ∼400 kb in the genome (Table2 and Figure1). Interestingly, canonical SurR binding motifs were detected only in the promoters of Mrp-mbh2 and SHII, while mrp-mbh1 and SHI contained several mutated motifs (Table2). Moreover, these gene clusters (*mrp-mbh1* and *shI*) showed consistently higher expression than their respective duplicates. For the sulfane sulfur reductase complex, *mrp-mbs*, four SurR binding sites were identified, with half exhibiting mutations. Additionally *surR* and *pdo*/*glutaredoxin* share two binding sites in their promoter region (Table2 and Figure 1). *T. barophilus* also possesses a unique Mrp-Mbh-Codh complex, enabling growth on carbon monoxide (Kozhevnikova et al., 2016). While its expression is regulated by pressure variation (Vannier et al., 2015), it has been proposed that in *Thermococcus onnurineus,* this complex is controlled by a regulator other than SurR (Lee et al., 2016). In *T. barophilus*, despite the presence of seven short binding sites within its cluster, only one mutated binding site is found in the promoter of the first coding sequence (Table 2). In this study, with the exception of Mrp-Mbh-Codh complex, all targeted ogeneclusters and SurR were genetically deleted. Their effects on physiological growth were assessed in sulfur-present and sulfur-absent conditions across different hydrostatic pressures (0.1, 40 and 70 MPa)

**Figure 1:** Genetic organization of gene clusters involved in hydrogenogenic/sulfidogenic metabolisms and their SurR binding motifs. The MBH locus (A) composed of two Mrp-Mbh clusters, the MBS locus (B), the SHII locus (C), the SHI locus (D) and the SurR/Pdo locus (E).

### Growth of wild type and derivative mutants under sulfur and hydrostatic pressure variations

Since genetic modifications were performed at atmospheric pressure (0.1 MPa), we first assessed the physiological impact of mutations under these conditions (Figure 2). Growth was evaluated with (Figure 2A) or without sulfur (Figure 2B). With sulfur, the Δ*mbh* strain exhibited only a slight growth delay compared to WT (see Table 3 for growth rate) while Δ*mbs* showed a more pronounced delay and reduced growth rate, leading to lower biomass after 24 h. Without sulfur, both Δ*mbh* and Δ*mbs* experienced severe growth impairment, failing to exceed a 1-log increase in cell density within 24 h, with growth rates below 0.1 h⁻¹ (0.023 and 0.032, respectively). This contrasts with previous findings in *P. furiosus* and *T. kodakarensis*, where the Mrp-Mbs cluster had no impact on sulfur-free growth (Kanai et al., 2005;Bridger et al., 2011;Kanai et al., 2011;Santangelo et al., 2011). Growth experiments at 40 MPa, the optimal pressure for *T. barophilus* (Figures 2C and 2D), showed that in the presence of sulfur, both mutants experienced initial growth delays but reached WT-like final densities after 24 h. Without sulfur, delays were more pronounced, and final cell concentrations were significantly lower than WT (Figure 2D, Table 3). Notably, despite lower overall growth rates, Δ*mbh* initially compensated for its delay after 15 h, while Δ*mbs* did so after 24 h, suggesting cooperative roles for MBS and MBH in sulfur-independent growth. This differs from previous findings, where only one of these clusters was necessary depending on sulfur availability. Additionally, the overexpression of mrp-mbh clusters at 0.1 MPa in sulfur-rich conditions (Vannier et al., 2015) does not appear essential for survival, as their deletion did not impact growth.

**Figure 2:** Characterization of Δ517, Δ*mbh* and Δ*mbs* mutants at 0.1 MPa (A & B) and 40 MPa (C & D). Growth assays were carried out in TRMm medium at 85°C, with sulfur (A & C) or without sulfur (B & D).

Next, we assessed strains lacking cytosolic hydrogenases under sulfur conditions at 0.1 MPa. With sulfur, Δ*shI* exhibited a more pronounced log-phase delay than ΔshII (Figure S2A). Without sulfur, Δ*shI* displayed the most significant growth impairment, reaching lower cell concentrations than WT and Δ*shII* after 24 h (Figure S2B). These results suggest that some sulfur-grown mutants can metabolically adapt, but those lacking membrane-bound clusters (Δ*mbh*, Δ*mbs*) cannot, likely due to the combined effects of genetic background and sulfur-dependent metabolic constraints.

Finally, we examined SurR’s role in growth across varying pressures and sulfur conditions (Figure 3, Table 4). Under sulfur conditions, Δ*surR* exhibited a growth delay at all pressures (Figure 3A). At 40 MPa, this delay was nearly compensated within 8 h, whereas at 70 MPa it took 10 h, and at 0.1 MPa over 14 h. Without sulfur, growth was severely impaired at all pressures (Figure 3B), but inhibition lessened as pressure increased, suggesting that pressure promotes cell recovery. Unlike non-piezophilic *Thermococcales* (e.g., *P. furiosus* and *T. kodakarensis*), where SurR deletion mainly affects sulfur-free growth (Santangelo et al., 2011;Lipscomb et al., 2017), our results indicate a role for SurR even in sulfur-containing conditions. This supports the hypothesis that SurR regulates hydrogen metabolism genes regardless of sulfur availability, potentially explaining their upregulation under sub-optimal pressure in piezophiles like *T. barophilus* (Vannier et al., 2015) and *T. piezophilus* (Moalic et al., 2021). Taken together, these findings highlight how pressure modulates molecular responses for cellular adaptation.

**Figure 3:** Characterization of Δ517 and Δ*surR* mutants at 0.1 MPa (A & D), 40 MPa (B & E) and 70 MPa (C & F). Growth assays were carried out in TRMm medium at 85°C, with sulfur (A & C) or without sulfur (B & D).

### Influence of SurR on the expression of targeted energetic metabolism genes

Given the distribution of canonical and variant SurR Binding sites in the promoters of genes and gene clusters involved in energy metabolism (Figure 1 and Table 2), we performed real-time quantitative PCR to assess SurR’s regulatory impact under varying sulfur and pressure conditions. Among the three reference genes tested, only pcna showed stable Ct values across experiments, and ΔCt calculations were based on this reference. Six target genes were selected from clusters encoding SHI, SHII, Mrp-Mbh1, Mrp-Mbh2, MBS, and SurR, based on their expression levels in sulfur conditions (Vannier et al., 2015). Since hydrogenogenic gene cluster expression increases at 0.1 MPa compared to 40 MPa in sulfur conditions, we compared their expression in the WT (Δ517) and SurR mutant strain (Δ*surR*) under three pressures (0.1, 40, and 70 MPa) with and without sulfur (Figure 4). At 0.1 MPa, in the presence of sulfur, ΔsurR exhibited significantly reduced expression for all target genes except *mbs* and *surR*, confirming SurR’s role in stimulating gene expression under suboptimal pressure despite sulfur availability (Figure 4A, left panel). Without sulfur, expression remained lower in Δ*surR*, though standard deviations overlapped, except for mbhI and shI (Figure 4A, right panel). A At 40 MPa, in the presence of sulfur, all genes were downregulated in Δ*surR*, though standard deviations overlapped for *mbs* and *surR* (Figure 4B, left panel), reinforcing SurR’s role in regulating hydrogenogenic genes under sulfur conditions. Without sulfur, all genes were downregulated except mbs and surR, which were overexpressed, suggesting distinct regulatory mechanisms (Figure 4B, right panel). At 70 MPa, a distinct regulatory pattern emerged, with shII upregulated 18-fold and mbhII threefold (Figure 4C, left panel). Without sulfur, expression patterns resembled those at 40 MPa, but with stronger *mbs* upregulation (Figure 4C, right panel). These findings confirm SurR’s influence on hydrogen metabolism gene regulation, particularly for *mbhI* and *shI*, under suboptimal and optimal pressure conditions. At 70 MPa, SHII and Mrp-Mbh2 appear to play a greater role, suggesting specific associations between membrane-bound complexes and cytosolic hydrogenases: SHII with Mrp-Mbh2, and SHI with Mrp-Mbh1. This aligns with the similarity of their SurR binding motifs (Figure 1, Table 2)

**Figure 4:** Effect of SurR DNA binding domain deletion on Gene expression variation of genes involved in hydrogenogenic/sulfidogenic metabolisms. The bars represent the expression ratio relative to PCNA gene (proliferating cell nuclear antigen). The blue bars symbolize the expression ratio of the Δ517 strain while the orange bars symbolize the expression ratio of the Δ*surR* strain. Growth rates are indicated in Table 4

To further investigate sulfur-driven regulation, we measured gene expression 30 minutes after sulfur addition (0.5 g/L or 2 g/L). As 0.25 g/L supports normal *T. barophilus* growth, these concentrations were chosen to enhance regulatory signals. In the WT, sulfur addition strongly repressed hydrogenogenic gene expression (Figure 5, upper panels), particularly *mbhI* and *shI*, whereas *mbs* and *surR* showed inconsistent trends between sulfur concentrations. In Δs*urR,* at 0.5 g/L, expression responses were more variable, with *mbhI* and *shII* showing strong fluctuations, while *mbhII* and *shI* decreased clearly (Figure 5, lower left panel). At 2 g/L, all hydrogenogenic genes except *shI* were downregulated, while *mbs* and *surR* displayed high variability (Figure 5, lower right panel). These results, obtained at 0.1 MPa, confirm that sulfur addition triggers a metabolic shift, reducing hydrogenogenic gene expression. They also reinforce SurR’s role in sulfur regulation, as repression was less pronounced in Δ*surR*, consistent with findings *in P. furiosus* (Lipscomb et al., 2017).

## CONCLUSION

This study provides new insights into the regulatory mechanisms controlling energy metabolism in *Thermococcus barophilus* under varying pressure and sulfur conditions. We demonstrat, that SurR, a redox-sensitive transcriptional regulator, plays a central role in metabolic adaptation, balancing hydrogen and sulfur metabolism in response to pressure. Under suboptimal pressure (0.1 MPa), SurR activates hydrogenogenic genes despite sulfur availability. At optimal (40 mPa) and high pressure (70 MPa), SurR’s regulatory dynamics shift, leading to a distinct reconfiguration of gene expression.

We also identify a functional association between membrane-bound hydrogenases (Mrp-Mbh1/Mrp-Mbh2) and cytosolic hydrogenases (SHI/SHII), in the adaptive response to pressure. Given that pressure affects protein folding activity (Chen and Makhatadze, 2017), further investigation into the three-dimensional structure of SurR and its physical interaction with DNA promoters under these pressure conditions is needed. Techniques such as real-time In Vitro Fluoresence anisotropy (Heisler et al., 2021) or Fluorescence Cross-Correlation Spectroscopy (FCCS), which can be applied under pressure (Bourges et al., 2017) provide valuable insights. Finally, integrating global “omic” approaches such as RNA-seq and Chip-seq will offer a comprehensive view of SurR regulation, helping to link its position at promoter regions to its role in gene regulation. These findings contribute to the broader field of extremophile biology and have implications for biotechnological applications and astrobiology, where life must adapt to fluctuating energy landscapes in high-pressure environments.

## DATA AVAILABILITY

The sequencing data were submitted to NCBI under BioProject accession number PRJNA1234085

## ACKNOWLEDGMENTS

We sincerely thank Marion Gardette, Betty Emonot, and Fatima-Zohra Masmoudi for their invaluable assistance in constructing and screening of the Δ*mbs* and Δ*surR* strains. We also extend our gratitude to Nadège Quintin for providing the strains used in this study, which are deposited in the UBO culture collection (https://ent.univ-brest.fr/lm2e/home/#/).

This work was supported by the Agence Nationale de la Recherche (ANR-10-BLAN-1725 01-Living deep; ANR-16-CE12–0016 CASPAR and ANR-22-CE02-0019-01 Hot Dog). A.T. received a postdoctoral fellowship from the Conseil départemental du Finistère and Ifremer. Y. M and T.B were supported by UBO

## FIGURE LEGENDS

**Figure S1:** Characterization of Δ517 strains with or without sulfur at 0.1 MPa. Growth assay were carried out at 85°C in TRM and TRMm media. TRMm containing Pyruvate (5g.L-1) was used for Δ517p strain growths. The Δ517p strain was previously adapted 96h in this medium before being subcultured for the growth kinetics. The growth rates are 0.059 h^-1^ (Δ517, blue curve), 0.35 h^-1^ (Δ517 + S, orange curve), 0.41 h^-1^ (Δ517p, green curve) and 0.48 h^-1^ (Δ517p + S, red curve)

**Figure S2:** Characterization of Δ*shI* and Δ*shII* mutants at 0.1 MPa. Growth assays were carried out in TRMm medium at 85°C, with sulfur (A) or without Sulfur (B). With sulfur, their respective growth rates are 0.41 h^-1^ and 0.28 h^-1^, while without sulfur, their respective growth rates are 0.33 h^-1^ and 0.11 h^-1^

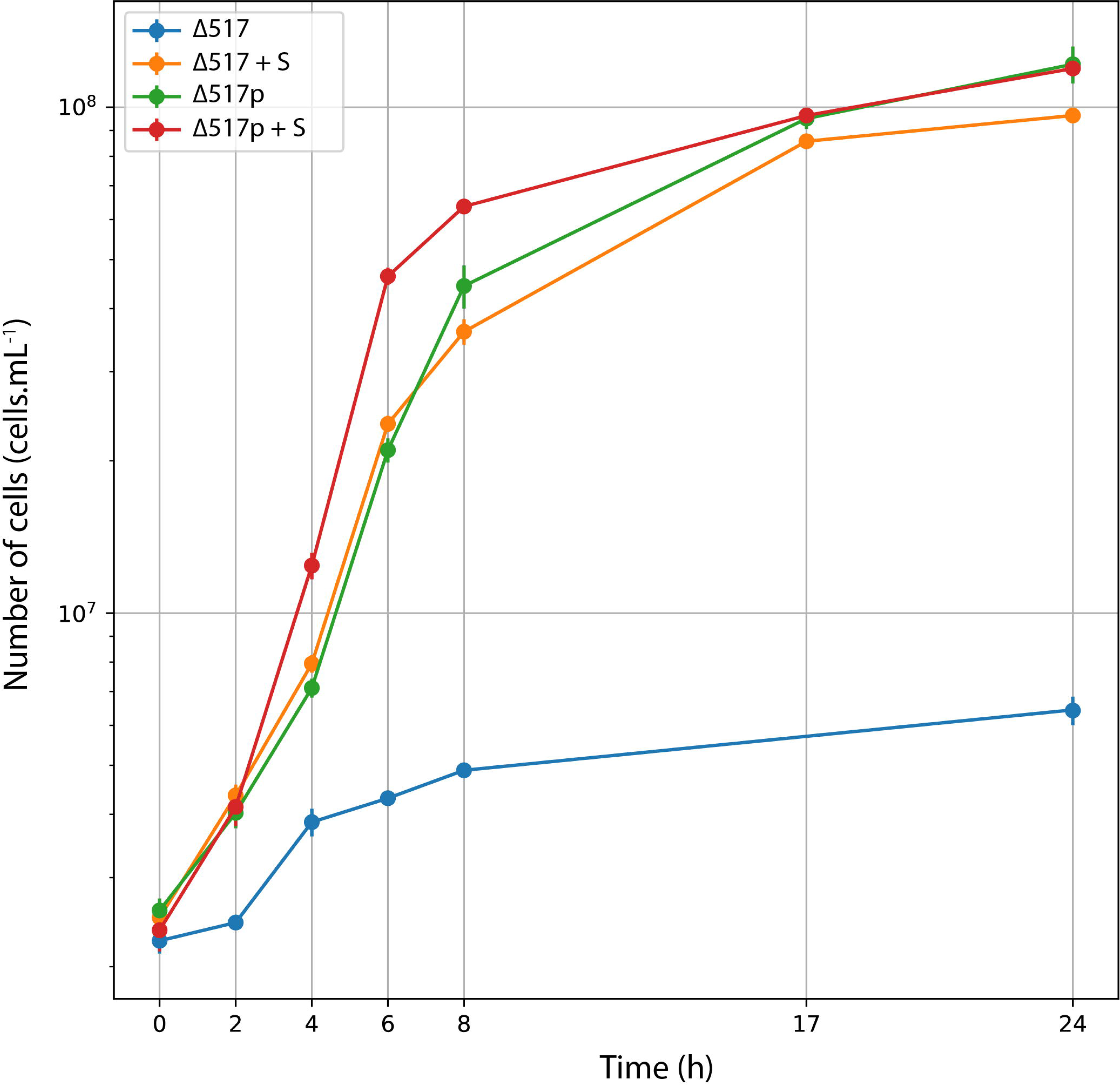

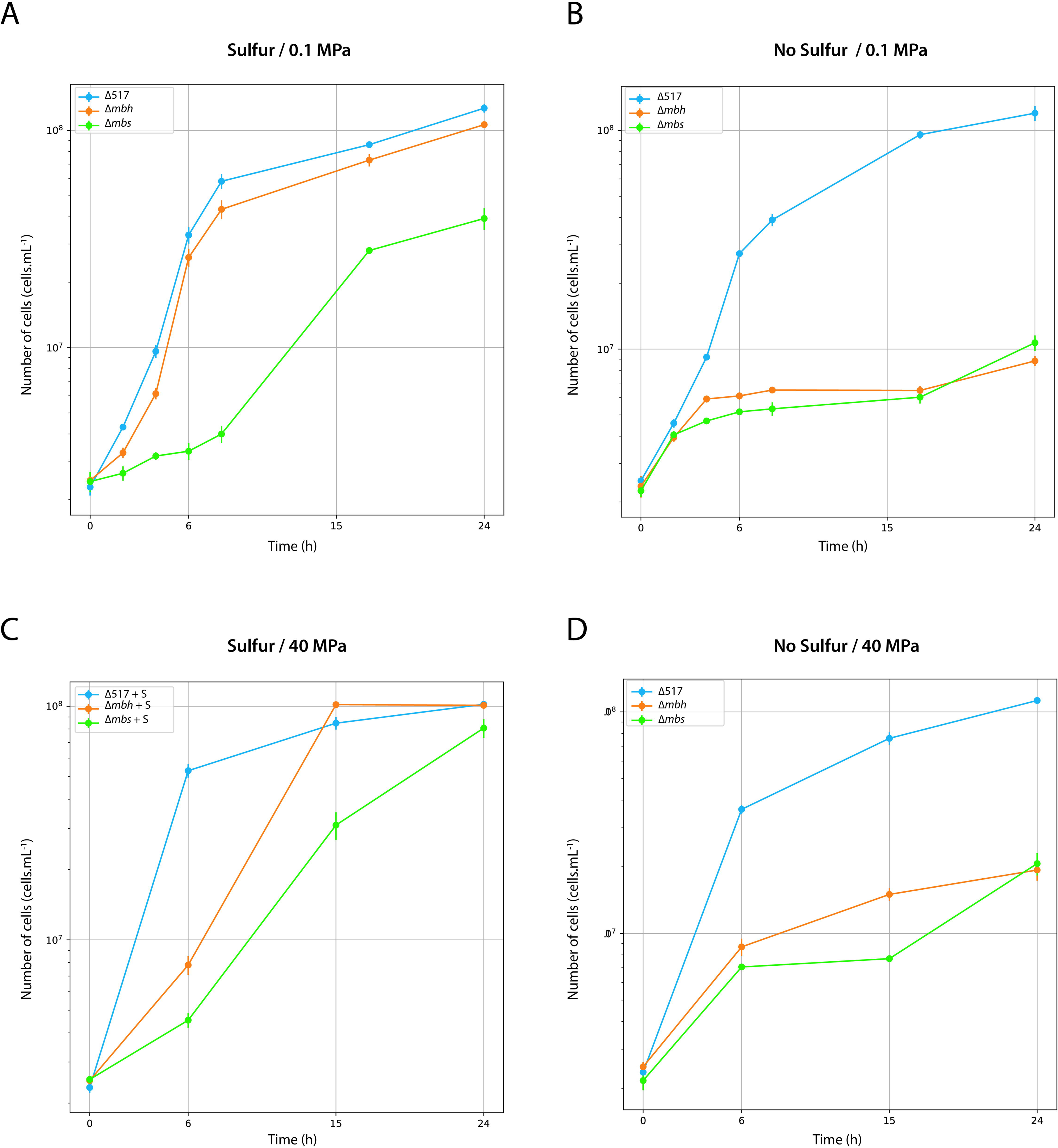

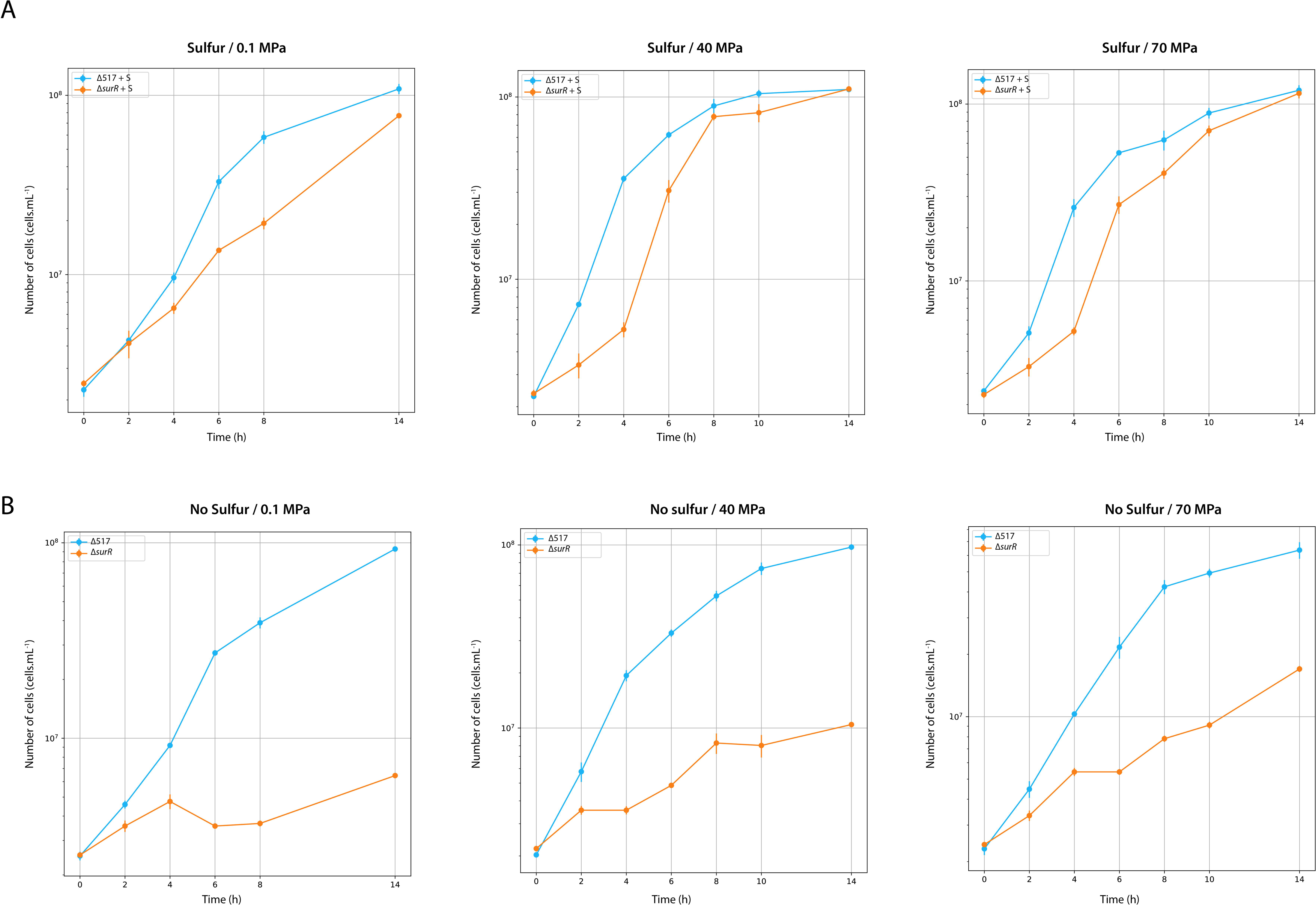

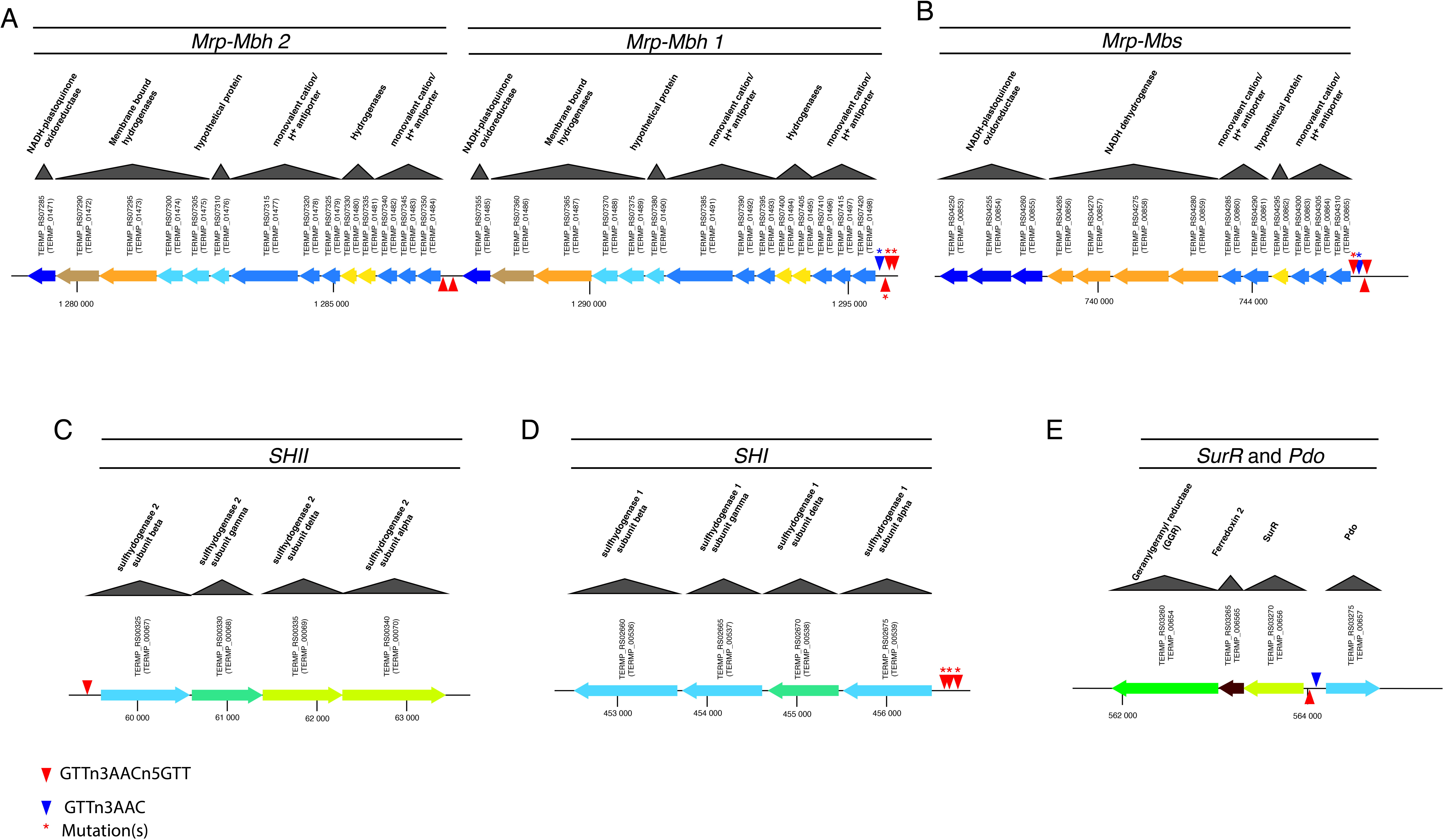

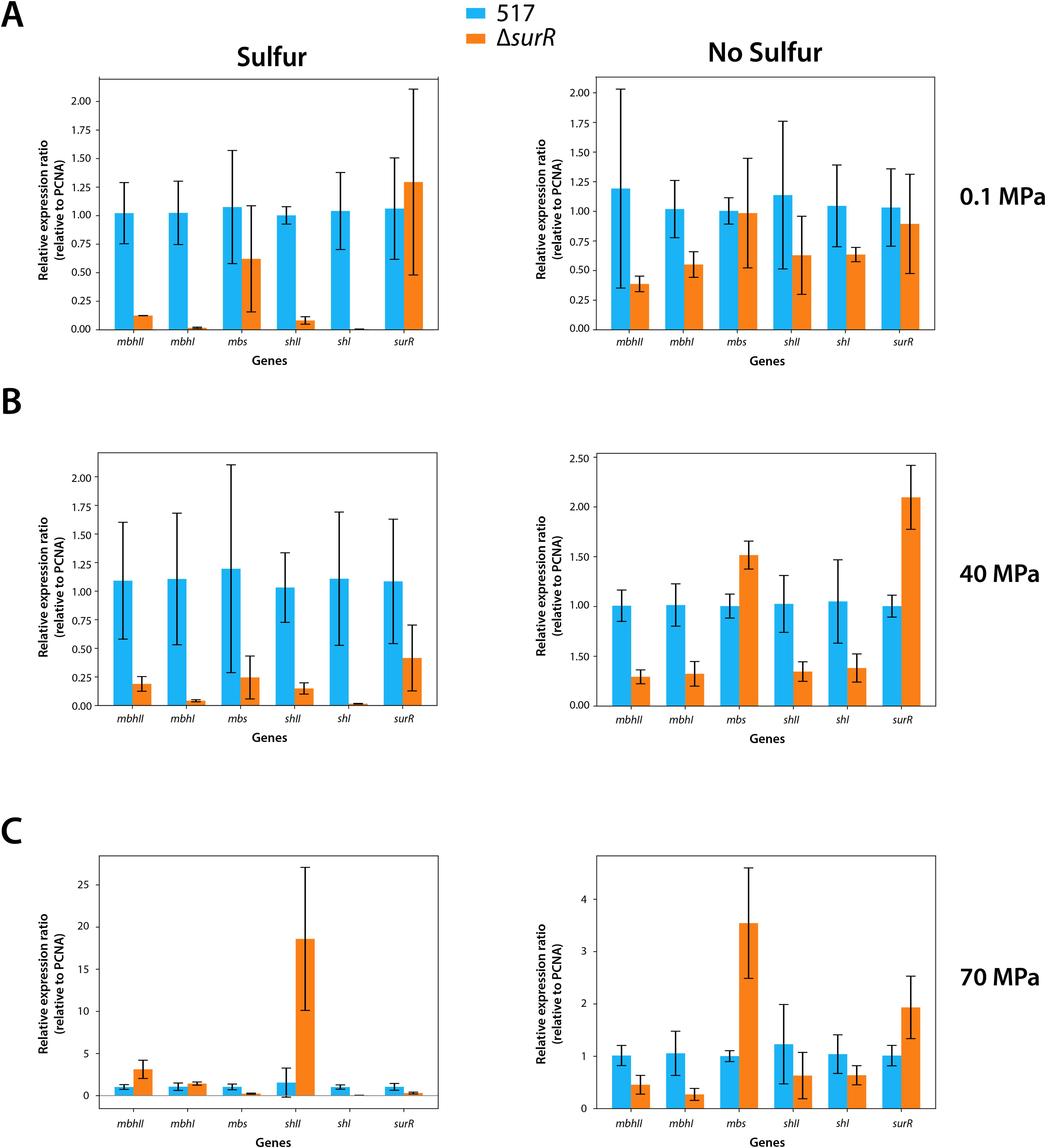

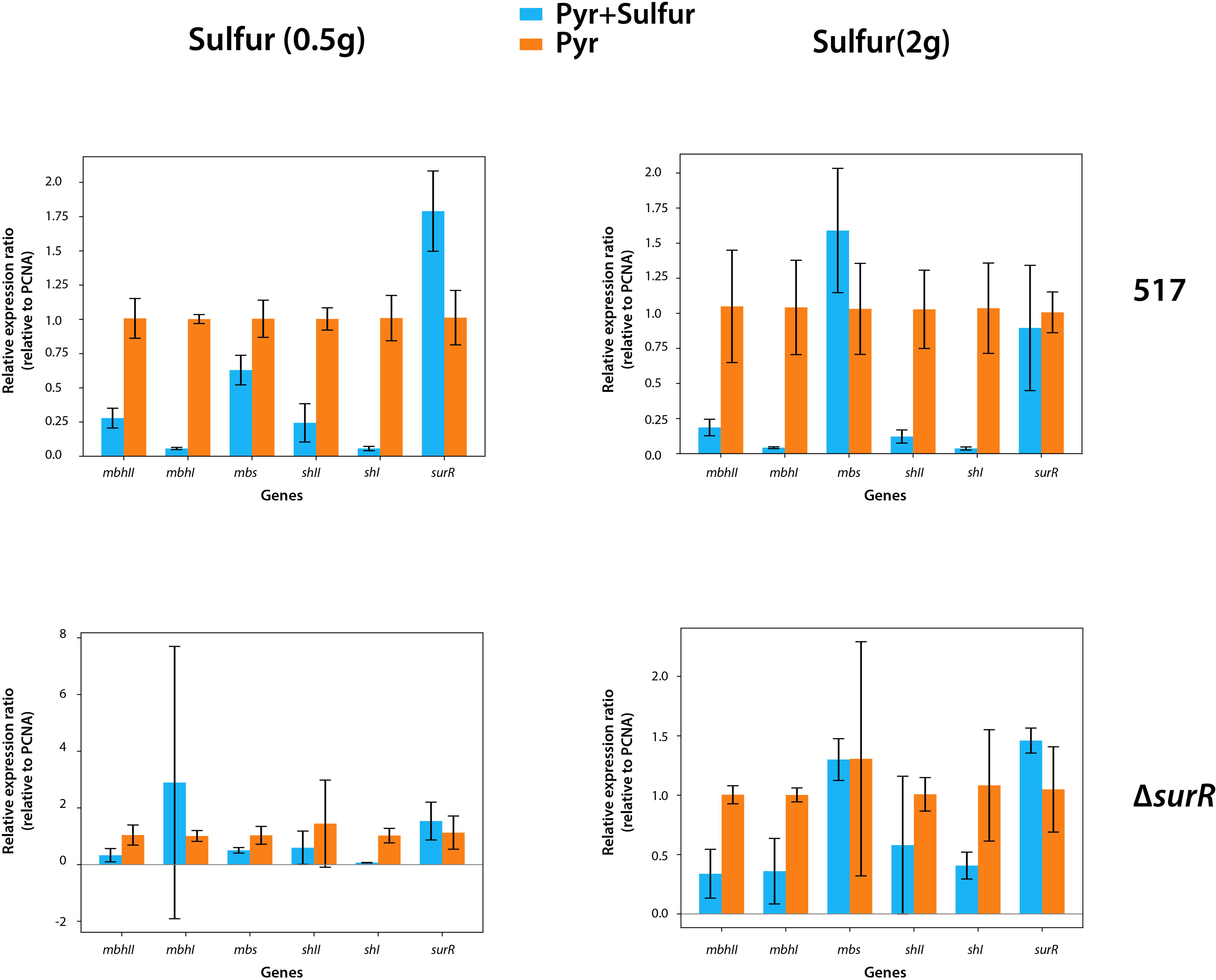

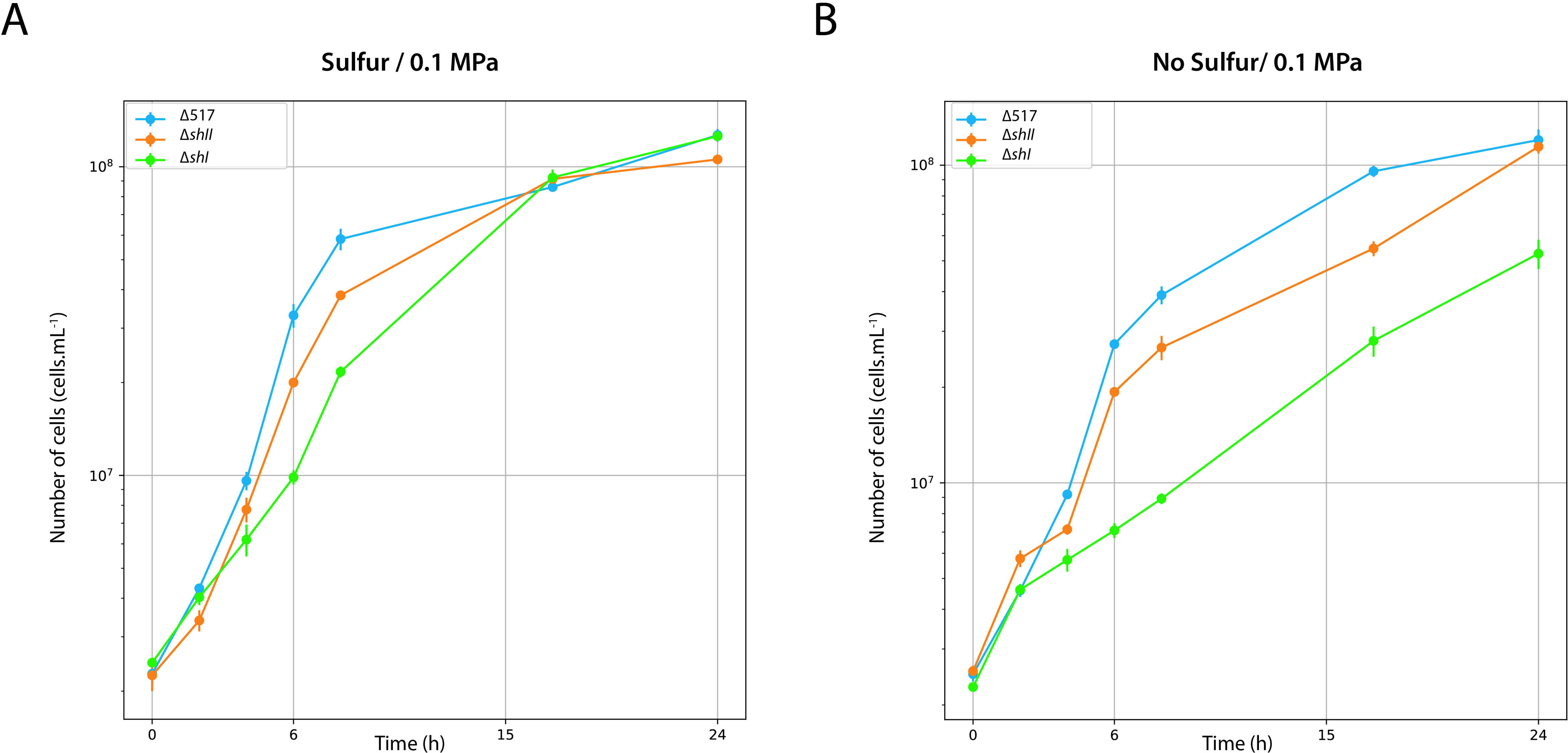

