## Supplementary material for "Regulation of gene expression under high hydrostatic pressure: the versatile role of the master regulator SurR in energy metabolism": Tables+Supplemental_Tables

Table1 : Specific Variants detected in the ∆517p strain and their potential effects

| **Genomic coordinate** | **locus** | **Uniprot ID** | **Protein Name** | **annotation impact** | **impact** |
| --- | --- | --- | --- | --- | --- |
| 434937 | TERMP_RS02585/TERMP_00520 | F0LJW0 | uncharacterized protein | frameshit | HIGH |
| 746729 | TERMP_RS04320/TERMP_00866 (upstream MBS)(GLY->ARG) | F0LLR6 | uncharacterized protein | missense_variant | MODERATE |
| 1348601 | TERMP_RS07735/TERMP_01561 THR->MET | F0LIT4 | low-affinity inorganic phosphate transporter | missense_variant | MODERATE |
| 1848338 | TERMP_RS10225/TERMP_02062 | F0LLK7 | Adenylyl-sulfate kinase (EC 2.7.1.25) | frameshit | HIGH |
| 1848341 | TERMP_RS10225/TERMP_02062 | F0LLK7 | Adenylyl-sulfate kinase (EC 2.7.1.25) | frameshit | HIGH |
| 1848405 | TERMP_RS10230/TERMP_02063 promoter | F0LLK8 | transposase | upstream_gene_variant | MODIFIER |

POS = genomic coordinates, Impact = Assessment of the putative impact of the variants through SnpEff.

Table2: SurR binding motifs positions upstream from the start codon of CDS involved in the electron flow

| **Product** | **Locus Tag**  **(read direction)** | **First CDS of the product** | **Motif occurrence** | **Upstream distance (size) of the Binding Motif from the start codon of the first CDS** | | |
| --- | --- | --- | --- | --- | --- | --- |
|  |  |  |  | **Upstream distance** | **Motif size** | **strand** |
| **Mrp-Mbh 1** | TERMP_RS07420-07355  (reverse) | TERMP_RS07420 | 4 | 21bp | Short with mutation: ATTn3AAC | forward |
|  |  |  |  | 119bp | Long with mutations: GTAn3AACn5TTT | forward |
|  |  |  |  | 139bp | Long with mutation: GTTn3AATn5GTT | reverse |
|  |  |  |  | 147bp | Long with mutation: ATTn3AACn5GTT | forward |
| **Mrp-Mbh 2** | TERMP_RS07350-07285  (reverse) | TERMP_RS07350 | 2 | 11bp | Long | reverse |
|  |  |  |  | 96bp | Long | reverse |
| **Mrp-Mbh-Codh** | TERMP_RS05755-05680 (reverse) | TERMP_RS05755 | 1 | 2bp | Short with mutation : GTTn3ACC | forward |
| **SHII** | TERMP_RS00325-00340  (forward) | TERMP_RS00325  (59 761 - 60 765) | 1 | 67bp | Long | forward |
| **SHI** | TERMP_RS02660-02675  (reverse) | TERMP_RS02675  (455 469 – 456 572) | 3 | 94bp | Long with mutation: GTTn3AAGn5GTT | reverse |
|  |  |  |  | 116bp | Long with mutations: GTTn3ATGn5GTT | forward |
|  |  |  |  | 257bp | Long with mutation: GTTn3AAAn5GTT | forward |
| **Mrp-Mbs** | TERMP_RS04310-04250  (reverse) | TERMP_RS04310 | 4 | 17bp | Long with mutation: GTTn2AACn6GTT | forward |
|  |  |  |  | 53bp | Short with mutation: GTTn3AAT | forward |
|  |  |  |  | 87bp | Long | reverse |
|  |  |  |  | 95bp | Long | forward |
| **SurR** | TERMP_RS03270  (reverse) | TERMP_RS03270  (563276 – 563 965) | 2 | 52bp | Long | reverse |
|  |  |  |  | 108bp | Short | forward/reverse |
| **Pdo /Glutaredoxin** | TERMP_RS03275  (forward) | TERMP_RS03275  (564 112- 564 786) | 2 | 31bp | Short | forward/reverse |
|  |  |  |  | 80bp | Long | reverse |
| **short : GTT*n*3AAC, long:GTT*n*3AAC*n*5GTT** |  |  |  |  |  |  |

Table 3: Growth rate of ∆517, ∆*mbh* and ∆*mbs* mutants according to Sulfur and HHP conditions. Growth were realized in TRMm at 85°C. Pyruvate added in Sulfur absence conditions

|  | 0.1 MPa | | 40 MPa | |
| --- | --- | --- | --- | --- |
| Strains | Sulfur | No Sulfur | Sulfur | No Sulfur |
| ∆517 | 0.45 h^-1^ | 0.38 h^-1^ | 0.51 h^-1^ | 0.45 h^-1^ |
| ∆*mbh* | 0.45 h^-1^ | 0.023 h^-1^ | 0.19 h^1^ | 0.21 h^-1^ |
| ∆*mbs* | 0.065 h^-1^ | 0.032 h^-1^ | 0.10 h^-1^ | 0.20 h^-1^ |

Table 4: Growth rate of ∆517 and ∆surR mutant according to Sulfur and HHP conditions. Growth were realized in TRMm at 85°C. Pyruvate added in Sulfur absence conditions

|  | 0.1 MPa | | 40 MPa | | 70 MPa | |
| --- | --- | --- | --- | --- | --- | --- |
| Strains | Sulfur | No Sulfur | Sulfur | No Sulfur | Sulfur | No Sulfur |
| ∆517 | 0.43 h^-1^ | 0.36 h^-1^ | 0.58 h^-1^ | 0.41 h^-1^ | 0.44 h^-1^ | 0.37 h^-1^ |
| ∆*surR* | 0.21 h^-1^ | 0.0041 h^-1^ | 0.56 h^-1^ | 0.10 h^-1^ | 0.38 h^-1^ | 0.13 h^-1^ |

TableS1: List of Strains used in this study

| Strains Names | UBOCC number | Genotype or other relevant characteristics | Genome region(s) deleted from parent strain | Source or reference |
| --- | --- | --- | --- | --- |
| ***E. coli*** |  |  |  |  |
| DH5α |  | *Φ80dlacZ*Δ*m15, recA1, endA1, gyrA96, thi-1, hsdR17 (r_k_−, m_k_+), supE44, relA1, deoR,* Δ*(lacZYA-argF)U169* |  | Thermo Fisher Scientific, Asnières, France |
| ***T. barophilus*** |  |  |  |  |
| ∆517 | UBOCC-M3300 | Δ*TERMP_00517* in TRM + Sulfur |  | Birien et al., 2018 |
| ∆*mbh* | UBOCC-M3309 | Δ*TERMP_00517* Δ*MBH* in TRM + Sulfur | From TERMP_1471 to TERMP_1498 | This study |
| ∆*shII* | UBOCC-M3311 | Δ*TERMP_00517* Δ*SHI* in TRM + Sulfur | From TERMP_00067 to TERMP_00070 | This study |
| ∆*shI* | UBOCC-M3312 | Δ*TERMP_00517* Δ*SHI* in TRM + Sulfur | From TERMP_00536 to TERMP_00539 | This study |
| ∆*mbs* | UBOCC-M3313 | Δ*TERMP_00517* Δ*MBS* in TRM + Sulfur | From TERMP_00853 to TERMP_00865 | This study |
| ∆*surR* | UBOCC-M3533 | Δ*TERMP_00517* Δ*SurR* in TRM + Sulfur (uncompleted deletion) | First 137bp of TERMP_00656 (nucleotides 563,829 – 563,965) | This study |
| ∆517p | UBOCC-M3535 | Δ*TERMP_00517* in TRMm + Pyruvate |  | This study |
| **Plasmids** |  |  |  |  |
| pUPH |  | Pop-in Pop-out vector |  | Birien et al., 2018 |
| pUFH-2 |  | Cloning of homologous regions flanking *TERMP*_*00005* |  | Thiel et al., 2014 |
| pUFH-3 |  | pUFH-2 + *surR* |  | This study |
| p6MP-∆*mbh* |  | pUPH+ homologous regions flanking the  double clusters Mrp-Mbh1 and Mrp-Mbh2 |  | This study |
| p6MP-∆*mbs* |  | pUPH+ homologous regions flanking the   cluster Mrp-Mbs |  | This study |
| p6MP-∆*shI* |  | pUPH+ homologous regions flanking the cluster ShI |  | This study |
| p6MP-∆*shII* |  | pUPH+ homologous regions flanking the   cluster ShII |  | This study |
| p6MP-∆*surR* |  | pUPH+ homologous regions flancking the fisrt 137 bp of TERMP_00656 |  | This study |

Table S2: Primers used for genetic constructions

| Primers name | Sequences (5’->3’) | Utilization |
| --- | --- | --- |
| KpnI-DHyd-1up | AAAAAAGGTACCATAAGAATCTTGCCATAAGGGC | To delete the double cluster *Mrp-Mbh* |
| DHyd-1do | CAAATCAAGAGATGAGGTGAGAAAACTTTCCGCTTTAAGTTTCTTTTTATAATAA |  |
| DHYD-2-up | TTATTATAAAAAGAAACTTAAAGCGGAAAGTTTTCTCACCTCATCTCTTGATTTG |  |
| BglII-DHyd-2do | AAAAAAAGATCTTTTCTGCAAGCTCAATAGCTTC |  |
| Verif_Trans_Hyd_Up | TTTTCCTCCCCTGGAAACTTCTTCC | To analyze the deletion of *the double cluster Mrp-Mbh* |
| Verif_Trans_Hyd_Do | ATCGGAGGTGTAAGGAGAGATCTCAAG |  |
| KpnI-∆Mbs_1up | AAAAAAggtaccACGTCAACATTTATGAAGTAAAACGCTATCCT | To delete the cluster *Mrp-Mbs* |
| ∆Mbs_1do | CTTGGAGGGTTGGTATTATCTTTTTCTCTTTTCTTAGGCTTTTCTTACATGAGAAAAA |  |
| ∆Mbs_2up | AAGAGAAAAAGATAATACCAACCCTCCAAGAGTTTAAACTTGAAAG |  |
| BamHI_∆Mbs_2do | AAAAAAGGATCCGGGAAAGATTGAGCATTACTTTGATGAATATCCT |  |
| Verif_∆Mbs_YM_do | CTTCATATACCTGCACCTCAGCGT | To analyze the deletion of the cluster *Mrp-Mbs* |
| Verif_∆Mbs_YM_up | GCCGAGAGCTATTGGAATTCTGG |  |
| KpnI_∆SHII_1up | AAAAAAGGTACCGGCCTCTCTGGAGTTGGAACCCTCT | To delete the cluster *ShII* |
| ∆SHII_1do | GAGAAATTTTGAATAGACGGGATCATCACCCGAAAGATTGCTTCTC |  |
| ∆SHII_2up | ATCTTTCGGGTGATGATCCCGTCTATTCAAAATTTCTCTATCTATTTTTGTTAAATC |  |
| BamHI_∆SHII_2do | AAAAAAGGATCCTCCATTTTGTTCTCGAGCATTAAAGCAACA |  |
| Verif_trans_SHII_up | TTCTGCAAGAGCTTAGGAACGCA | To analyze the deletion of the cluster *ShII* |
| Verif_trans_SHII_do | TATGCATCCGAGTTTCGATTTAACACATATAGC |  |
| KpnI_DSHI_1up | AAAAAAGGTACCCCCAATAAAATAAGCTCATTTATGGCAGCG | To delete the cluster *ShI* |
| DSHI_1do | GGGGAGGGGATCTCTATTCTCTTTTCTAAATTTTATCTCTGGTGGTTTTATGA |  |
| DSHI_2up | TTTAGAAAAGAGAATAGAGATCCCCTCCCCATGAACATCAT |  |
| BglII_DSHI_1do | AAAAAAAGATCTAAAGACATTTTCGAGCTCCTCATACCC |  |
| Verif_SHI_up | CCGGGCTCAAATTTTTCACTTAGAATCG | To analyze the deletion of the cluster *ShI* |
| Verif_SHI_do | CAATGAACATCATTTGTCCTTCGCCAC |  |
| KpnI_∆SurR_YM4-1up | AAAAAAGGTACCGTCTATAGCCGCTCGGGGAATATATTG | To delete partially the gene *SurR* |
| ∆SurR_YM4-1do | TGAGTAGGTGGGGGACCCTCAAAATCATGGAAAGGGAGGGAC |  |
| ∆SurR_YM4-2up | CCCTTTCCATGATTTTGAGGGTCCCCCACCTACTCAAATTTGTAGC |  |
| BamHI_∆SurR_YM4-2do | AAAAAAGGATCCTAACATTTTTGCGAAAGCCATGGGAG |  |
| BamHI_SurR_int_up | AAAAAAGGATCCAACGGTACAAACTTGTTTAGCTCTTC | To analyze the partial deletion of the gene *SurR* |
| KpnI_SurR_int_do | AAAAAAGGTACCCATTCTGGGAAATAAGGTGAGGAGAGA |  |

Table S3: Primers used for the RT-QPCR

| **Targeted Gene/Cluster** | **Targeted Locus** | **Forward Primer (5’ to 3’)** | **Reverse Primer (5’ to 3’)** | **Expected Amplicon size (bp)** |
| --- | --- | --- | --- | --- |
| *sh*II | TERMP_00068 | TACCTACCGGCCAGGTCAGT | GGTCCCCGAA TGCCCACA | 180 |
| *sh*I | TERMP_00536 | GCATAGCTGACCC TTCCATTCTT | AACAACGCTGATCTGCTCTAT GG | 284 |
| *mbh*2 | TERMP_01494 | AGTGGCTTCTCCAAGGG TATCATAAC | ATCCTTGGTGTGCTCTTAATA GTAGCTAACC | 195 |
| *mbh*1 | TERMP_01480 | GTTCACGCTCTTCCTCCTTG | GTCGCTTCGCCAAGAGTATC | 174 |
| *mbs* | TERMP_00865 | GCAGCACTCTGTAG GCAACATC | TTCCTGACGAGCAGCCTTGAT C | 205 |
| *Sur*R | TERMP_00656 | CGTCCAGCTTTATGCCCCTGT   | CCTCAACGGCAGTCTCAAAAC A | 180 |
| *30S (S19)* | TERMP_00095 | AGTGGTGGCCGCTATTATTG | TAGGATTTCACCCCTACCCC | 156 |
| *30S (S13)* | TERMP_00128 | GGCATAAACTTTGCCACGATG GTG | TGCACCAATGAGGTGCATGTC | 195 |
| *pcna* | TERMP_00342 | GCATGAGGGCAATGGATCCA A | GGGTTACCTCAAGGAAGTTCT CCTCA | 198 |
